# Promoting neuronal outgrowth using ridged scaffolds coated with extracellular matrix proteins

**DOI:** 10.1101/788539

**Authors:** Ahad M. Siddiqui, Rosa Brunner, Gregory M. Harris, Alan.L. Miller, Brian E. Waletzki, Jean E. Schwarzbauer, Jeffrey Schwartz, Michael J. Yaszemski, Anthony J. Windebank, Nicolas N. Madigan

## Abstract

Spinal cord injury (SCI) results in cell death, demyelination, and axonal loss. The spinal cord has a limited ability to regenerate and current clinical therapies for SCI are not effective in helping promote neurologic recovery. We have developed a novel scaffold biomaterial that is fabricated from the biodegradable hydrogel oligo[poly(ethylene glycol)fumarate] (OPF). We have previously shown that positively charged OPF scaffolds (OPF+) in an open spaced, multichannel design can be loaded with Schwann cells to support axonal generation and functional recovery following SCI. We have now developed a hybrid OPF+ biomaterial that increases the surface area available for cell attachment and that contains an aligned microarchitecture and extracellular matrix (ECM) proteins to better support axonal regeneration. OPF+ was fabricated as 0.08 mm thick sheets containing 100 μm high polymer ridges that self-assembles into a spiral shape when hydrated. Laminin, fibronectin, or collagen I coating promoted neuron attachment and axonal outgrowth on the scaffold surface. In addition, the ridges aligned axons in a longitudinal bipolar orientation. Decreasing the space between the ridges increased the number of cells and neurites aligned in the direction of the ridge. Schwann cells seeded on laminin coated OPF+ sheets aligned along the ridges over a 6-day period and could myelinate dorsal root ganglion neurons over 4 weeks. The OPF+ sheets support axonal regeneration when implanted into the transected spinal cord. This novel scaffold design, with closer spaced ridges and Schwann cells is a novel biomaterial construct to promote regeneration after SCI.

**Graphical Abstract:** 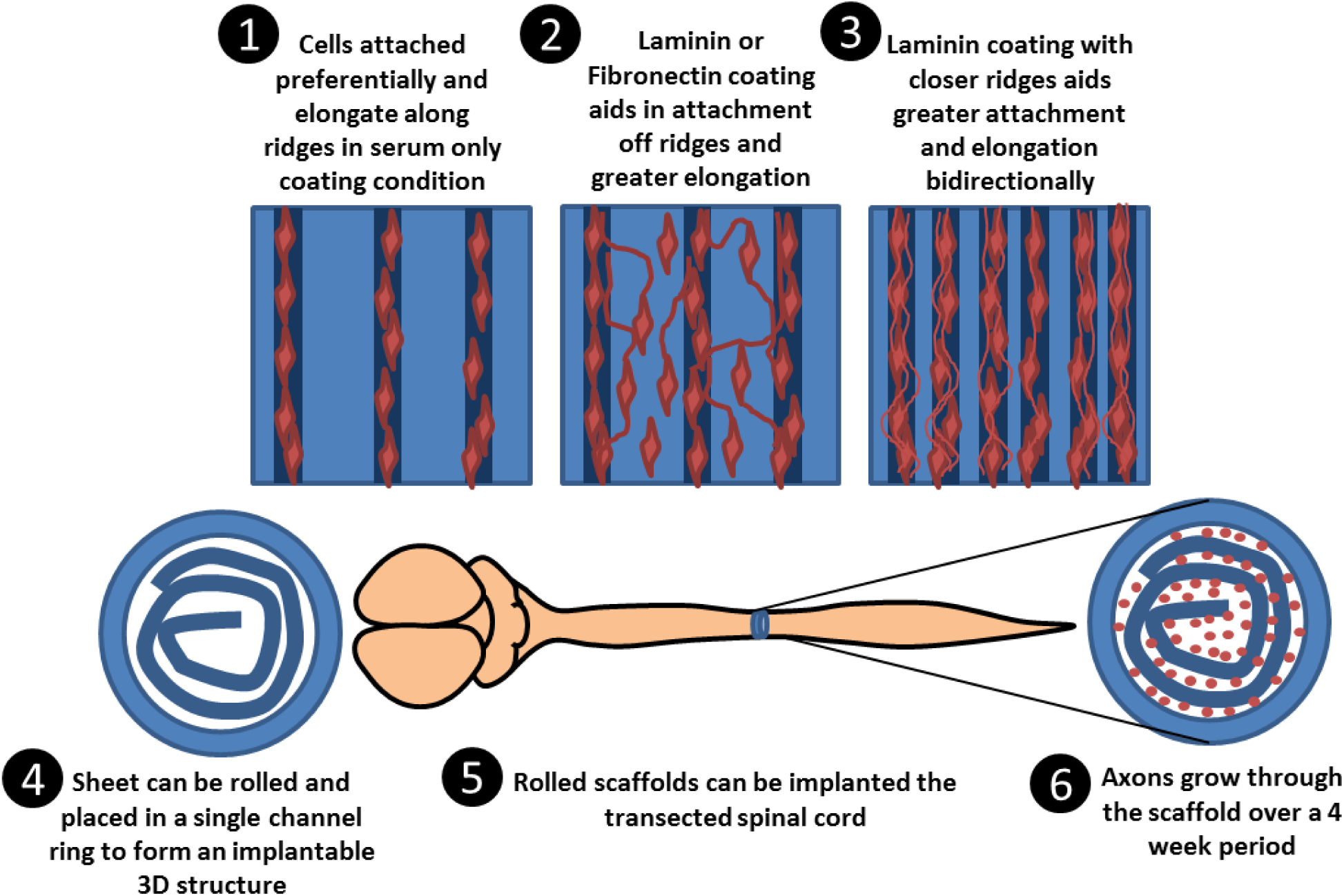

## 1. Introduction

The spinal cord has a limited ability to regenerate after spinal cord injury (SCI) and available therapies are not efficacious in promoting recovery of motor and sensory neurological function. The failure to recover function may be due to secondary events that occur after the primary insult such as cell death, axonal loss, demyelination, cyst formation and an increase in the inhibitory microenvironment [1]. Many therapies are currently under investigation for treating one or more of these secondary events using neuroprotective strategies and cell, regenerative, and rehabilitative therapies [1, 2]. A single therapeutic strategy is unlikely to be effective in treating a disorder that is heterogeneous in its underlying pathophysiology. Combinatorial treatments that simultaneously address multiple contributions to the residual of the injury will be required.

Biomaterial scaffolds are an attractive platform for combinatorial therapies because they are able to bridge the physical gap produced from gliosis, cyst formation, and cell death by providing structural support for regenerating cells [3, 4]. Biomaterials can be combined with extracellular matrices, cells, or pharmaceuticals to promote a more hospitable environment for regeneration [3, 5, 6]. We have developed a positively charged polymer, oligo[poly(ethylene glycol)fumarate] (OPF+) for scaffold fabrication in a multichannel design that overlays the different tracts of the spinal cord [4, 7]. OPF+ forms a porous, biodegradable hydrogel that is mechanically similar to the spinal cord [8]. The positive charge on OPF enhanced neuronal cell attachment, extension and the axonal myelination with Schwann cells *in vitro* [9]. *In vivo*, OPF+ loaded with Matrigel supported axonal regeneration over 4 weeks [10]. OPF+ scaffolds loaded with Schwann cells further improved axonal regeneration, and growth orientation over that seen with Matrigel alone, and reduced collagen scarring, cyst formation, and proteoglycan accumulation following transection injury [4, 7, 10]. Schwann cells, that were genetically modified to express high concentrations of glial cell-derived neurotrophic factor (GDNF-SC), increased axonal regeneration and myelination compared to wild type Schwann cells and led to enhanced functional recovery following spinal cord transection [5].

Axonal regeneration occurred through the open channels in the scaffold, as well as on the outer surface of the scaffolds. This observation led to the new hypothesis that changing the scaffold architecture to increase the surface area available for axons to regenerate may increase the number of axons regenerating through the scaffold and provide directional guidance. Different scaffold designs have been fabricated with a higher proportion of open to closed spaces in association with a novel ridged surface architecture that would guide regenerating axons. Physical structures for contact mediated axonal guidance, such as microgrooves or ridges, have been created by laser etching onto polymer surfaces [11-13]. Initial studies using quartz slides microgrooves improved the directional orientation of seeded fibroblast and epithelial cells but did not align chick embryo neurons [11]. Yao *et al.* (2009) subsequently demonstrated that micro-patterned poly(lactide-co-glycolide) (PLGA) sheets increased neurite alignment. In another study, neonatal rat dorsal root ganglion (DRG) neurons were cultured on grooved polymers of poly(dimethyl siloxane) coated with poly-L-lysine and laminin. On this surface neurites extended along and arched over the grooves with the somas preferentially adhering to the ridges [14].

The addition of extracellular matrix (ECM) proteins onto polymer surfaces provides sites for receptor-mediated cell attachment to promote axonal guidance and cell migration [15]. Grooved polymers coated with gradients of laminin or collagen have been shown to direct axonal growth cones [16, 17]. Fibronectin promotes Schwann cell proliferation and motility in culture [18], and is an attractive ECM candidate to make biomaterials more cell adhesive and facilitate growth. Peripheral nerve regeneration in rats can be enhanced by the addition of fibronectin and Schwann cells in alginate hydrogels [19]. Fibronectin has also been shown to help increase serotonergic axon sprouting and can influence axonal growth when used in combination with cells, growth factors, and biomaterials [20, 21].

In this study we sought to determine which ECM molecules best support neuronal attachment, and whether the structural microarchitecture may further improve the directional alignment of axonal extension and myelination *in vitro*. A novel hybrid OPF+ scaffold design was fabricated as a ridged, flat sheet and coated with ECM proteins. Upon hydration the sheets had the unique property of spontaneously rolling into a spiral 3D configuration. Polymer sheets with ridge spacings of 0.2 mm, 0.4 mm, or 1 mm apart were coated with laminin, collagen, or fibronectin, and used in *in vitro* assays to quantitate neuronal attachment, DRG neurite extension, and axonal myelination by Schwann cells. A proof of concept study was then conducted *in vivo* to compare axonal regeneration in laminin and untreated spiral scaffolds implanted following complete spinal cord transection.

## 2. Material and Methods

### 2.1 OPF Synthesis and OPF+ Scaffold Fabrication

OPF+ was synthesized as described previously [8, 9]. To create the ridged OPF+ sheet, liquid polymer was pipetted onto a Teflon mold that contained micro-grooves 100 µm in depth and spaced 1 mm, 0.4 mm and 0.2 mm apart. A glass slide was place on top with a 0.08 mm thick spacer. The sheets were polymerized by exposure to an ultra-violet (UV) light (365nm) for one hour and cured overnight. Single channel OPF+ scaffolds were fabricated by mold injection of the polymer, and cast over a 1mm wire prior to UV exposure. The scaffolds were cut into 2mm lengths for transplantation.

### 2.3 Swelling Ratio

Desiccated, ridged OPF+ sheets were cut into 6×6 mm pieces and weighed dry (W_d_) and following hydration in distilled water for 24 hours (W_s_; swollen weight).

The swelling ratio was calculated using the equation:

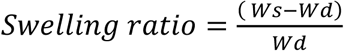

### 2.4 OPF+ Sheet Culture Preparation and ECM Protein Coating

OPF+ sheets were sterilized by immersion in serial dilution of ethanol, rehydrated in cell culture media and then pinned flat onto sterile culture dishes containing a layer of silicone elastomer (SYLGARD^®^ 184 Silicone Elastomer Kit; Dow Corning, Midland, MI, USA). The OPF+ sheets containing ridges spaced 1 mm apart were coated with rat tail collagen 1 (3.36 mg/ml; Corning^®^, New York, USA), laminin (100 μg/ml; Sigma-Aldrich^®^, St. Louis, MO, USA), or rat plasma fibronectin (10 μg/ml) overnight at 37°C. Control sheets received no additional ECM protein coating other than that which would be present in the serum of the media (referred to as serum coated). The fibronectin was purified by gelatin-Sepharose affinity chromatography from frozen rat plasma [6]. The sheets were then washed 3 times to remove unabsorbed ECM proteins. To determine the effect of dissociated DRG attachement and alignment on OPF+ sheets with different ridge sizes, OPF+ sheets with ridges 0.2 mm, 0.4mm, and 1mm apart were coated with laminin.

### 2.5 Whole DRG Explants

DRG neurons were isolated from Sprague Dawley pups on embryonic day 15 as previously described [22]. Whole DRG were placed on the scaffold in media containing modified Eagle’s medium (MEM; Gibco-BRL, Gaithersburg, MD) supplemented with 10% calf bovine serum (CBS) (HyClone, Logan, Utah, USA), L-glutamine (1.2 mM; Gibco), glucose (7mg/ml; Sigma Aldrich) and nerve growth factor (NGF, 5ng/ml; Harlan Bioproducts, Indianapolis, IN, USA). Initially, the explants were incubated for 1 hour at 37°C in 150 μL media to allow them time to attach. Once the DRG explants were attached the plate was filled with media.

Eighteen DRGs for each condition were plated with 4-6 DRGs per scaffold. DRG explants were imaged at 24 and 48 hours after isolation using a Zeiss Axiovert Model 35 microscope with a Nikon CCD camera. Length of the longest neurite of each DRG was analyzed by measuring from the edge of the DRG to the end of the longest outgrowing neurite using Image J (NIH). Given a variable degree of DRG attachment on each type of substrate, the final numbers of explants available for analysis was n=17 DRGs for laminin, n=11 DRGs for collagen, n=11 DRGs for fibronectin, and n=4 DRGs for serum coated.

### 2.6 DRG Neuronal Cultures

Dissociated DRG neurons were isolated from Sprague Dawley pups on embryonic day 15 as previously described [22]. Approximately 100-200 whole DRGs were pooled and dissociated using trypsin and mechanical trituration. 50,000 cells were plated onto each scaffold sheet, and were grown in MEM supplemented with 15% CBS (HyClone, Logan, Utah, USA), L-glutamine (1.2 mM; Gibco), glucose (7 mg/ml; Sigma Aldrich) and NGF (5ng/ml; Harlan Bioproducts, Indianapolis, IN, USA). Dissociated cultures were treated for 3 days with 1 x 10^−5^ M 5-fluoro-2-deoxy-uridine and with 1 x 10^−5^ M uridine (Sigma) to remove non-neuronal cells[23, 24]. The cells were fixed with 4% paraformaldehyde for 30 minutes and stored in phosphate buffered saline (PBS) at 4°C for immunocytochemistry.

### 2.7 Immunocytochemistry

The fixed OPF+ sheets were blocked with 10% normal donkey serum and 0.1% Triton X-100 in 0.01M PBS for 30 minutes. Primary antibodies against mouse anti-β-III tubulin (1:300, Millipore) and rabbit anti-myelin basic protein (1:800, Abcam) were diluted in PBS containing 5% normal donkey serum and 0.3% Triton X-100 at 4°C overnight. Secondary antibodies (donkey anti-mouse Cy3; Millpore; 1:200 and donkey anti-rabbit Alexa 647; Jackson ImmunoResearch Laboratories; 1:200) were diluted with 5% normal donkey serum and 0.3% Triton-X-100 in PBS. After removing the pins, the OPF+ sheets were carefully placed onto a glass slide using forceps with the ridges facing upwards. Glass coverslips were mounted using Slow Fade Gold Antifade Reagent with DAPI nuclear stain (Molecular Probes^©^, Eugene, Oregon, USA).

### 2.8 Image Analysis of DRG Neurons

All images of dissociated neurons were taken using an inverted fluorescence microscope Zeiss Axio Observer Z-1 with a motorized stage (Carl Zeiss, Inc., Oberkochen, Germany) mounted with an Axiocam 503 camera (Zeiss). Pictures were acquired via the ZEN 2 (blue edition) imaging software (Carl Zeiss). For cell quantification, z-stacks with approximately 5 slices at magnification of 10x were taken using an EC Plan-Neofluar 10x/0.30 Ph1 objective. On each polymer sheet, 4 representative areas of different ridges and the adjacent flat parts were systematically captured. To avoid bias, the regions were chosen by moving one field of view up or down form the centered position and then 2 fields across left or right to yield a total of 4 quadrants. The DRG neurons preparation was repeated at least 3 times. Cell nuclei (DAPI), which were associated with axons (β-tubulin, red) were counted in each image in an area of 878.94 µm x 662.84 µm on the polymer to quantify neuronal cell attachment.

In order to determine the ECM protein-coating that displayed most neurite outgrowth and attachment, the amount of β-III tubulin was calculated for each polymer sheet using ImageJ software by measuring the mean gray value of the staining. The amount of β-tubulin staining was normalized to the amount of DAPI (mean grey value) per cell. A mean value of normalized β-tubulin mean gray values was derived for each polymer sheet from replicates of the 4 fields of views. For comparison between different ECM molecules on OPF+ sheets with 1 mm spaced ridges, laminin coated sheets had n=7 sheets, fibronectin coated sheets had n=3 sheets, collagen coated sheets had n=4 sheet, and sheets with no substrate had n=6 sheets. For different spaced ridges, OPF+ sheets with 0.2 mm spaced ridges had n=3 sheets, 0.4 mm spaced ridged sheets had n=2 sheets, and 1 mm ridged sheets had n=4 sheets.

### 2.9 Schwann Cell Cultures

Primary rat Schwann cells were isolated from sciatic nerves of 2-5 day old Sprague Dawley pups as previously described [25, 26]. 125,000 cells were seeded on to laminin coated OPF+ ridged sheets. Schwann cells were imaged using Zeiss Axiovert Model 35 microscope with a Nikon CCD camera at 24 hours and 6 days. For each condition 3 scaffold replicates were made and images were taken of 4 fields of views per scaffold.

### 2.10 Neuronal-Schwann Cell Co-Culture

Schwann cells and dissociated DRG neurons were isolated and plated together (1:1 ratio, 100,000 cells in total per scaffold) on OPF+ ridged sheets. The cells were cultured for 4 weeks in MEM supplemented with 15% CBS (HyClone, Logan, Utah, USA), L-glutamine (1.2 mM; Gibco), glucose (7 mg/ml; Sigma Aldrich) and NGF (5 ng/ml; Harlan Bioproducts, Indianapolis, IN, USA), 50 μg/mL of ascorbic acid (Sigma) and 2 μM Forskolin (Sigma). For each condition 3 scaffold replicates were made and images were taken of 4 fields of views per scaffold using a Zeiss LSM780 confocal microscope system.

### 2.11 OPF+ sheet and scaffold preparation and surgical implantation

OPF+ sheets and single channel scaffolds were sterilized with serial dilution of ethanol. 2 mm by 6 mm OPF+ sheets with ridges 0.4 mm apart were either coated with laminin (n=1; 100 μg/ml) or had only serum coating (n=1). The scaffold sheet was rolled along the long edged and placed inside the single channel scaffold with 2 mm length. Female Sprague Dawley rats (230 – 300 grams; Harlan Laboratories, Madison, WI) received a laminectomy at level T8-T10 and a complete spinal cord transection was performed at level T9. The scaffold was placed into the transected area and all the muscles were closed with sutures. The animals were cared for by the research team and by veterinarians their staff experienced with care of rats with spinal cord injury daily. All procedures were approved by the Mayo Clinic Institutional Animal Care and Use Committee and all guidelines were followed in accordance with the National Institute of Health, Institute for Laboratory Animal Research and the United States Public Health Services Policy on the Humane Care and Use of Laboratory Animals.

### 2.12 Tissue preparation and Immunohistochemistry

The animals were euthanized 4 weeks after implantation and injury by deep anesthesia and fixed by transcardial perfusion with 4% paraformaldehyde in PBS. The spinal column was removed en bloc and post fixed for 2 day at 4°C. The spinal cords were dissected out and a 1 cm segment with scaffold (T9) in the center was embedded in paraffin. Transverse sections 8 μm thick were made on a Reichert-Jung Biocut microtome (Leica, Bannockburn, IL). Tissue sections were deparaffinized and immunostained as we have described (Chen et al 2017) using a mouse anti-human neurofilament primary antibody (1:100; DAKO) and a donkey anti-mouse IgG secondary antibody conjugated to Cy3.

### 2.13 Imaging for axon regeneration in scaffolds

The sections were imaged using Zeiss Axiovert Model 35 microscope with a Nikon CCD camera. The microscope contained a motorized stage and the tiling feature was used in the Zen Blue (Zeiss) software to image the whole scaffold section. Neurofilament staining identified the axons running through the scaffold.

### 2.14 Statistics

All data are reported as mean ± standard error of the mean (SEM). All statistics were calculated using GraphPad Prism 7 (GraphPad Software, Inc., USA). Statistical significance was calculated using one-way ANOVA with multiple comparisons using Tukey’s post-hoc testing (unless stated otherwise). On graphs p-values are represented as *p< 0.05, ** p< 0.01, *** p< 0.001, **** p< 0.0001.

## 3. Results

### 3.1 Ridged OPF+ Sheet Characterization

0.08 mm thick OPF+ sheets were embedded with longitudinal ridges spaced 0.2 mm, 0.4 mm, or 1 mm apart (Figure 1A). Our previously used scaffold was fabricated with 7 channels that were 2mm wide and 32 mm long (Figure 1B). Swelling ratios were calculated to be 10.5 ± 0.8 for OPF+ sheets with 1 mm spaced ridges, 9.8 ± 0.4 for sheets with 0.4 mm spaced ridges, and 9.7 ± 0.1 for sheets with 0.2 mm spaced ridges (Figure 1C). Hydrated sheets spontaneously rolled into a spiral configuration (Figure 1D and 1E). In comparison to a multichannel scaffold (Figure 1B), the rolled-sheet design allowed for a greater volume of open space inside of the scaffold, and increased surface area that was dependent on the number of loops in the spiral (Figure 1F and 1G). Internal volumes and surface areas were calculated by taking into account the number of channels that form between ridges, ridge separation (mm), length of the cylinder (mm), and ridge width (mm) (Figure 1F). A scaffold sheet with 3x loops increased the volume by 2.6 fold (3.34 mm^3^ to 8.83 mm^3^), whereas a sheet with 4x loops has an increase of approximately 3 fold (3.34 mm^3^ to 9.72 mm^3^) compared to the 7 multichannel OPF+ scaffold design (Figure 1F). There was a similar corresponding increase in surface area where sheets with 3x loops had 1.71 fold increase (45.4 mm^2^ to 77.7 mm^2^) and 4x loops had 2.87 fold increase (45.4 mm^2^ to 130.5 mm^2^) compared to the 7 channel design.

**Figure 1:**
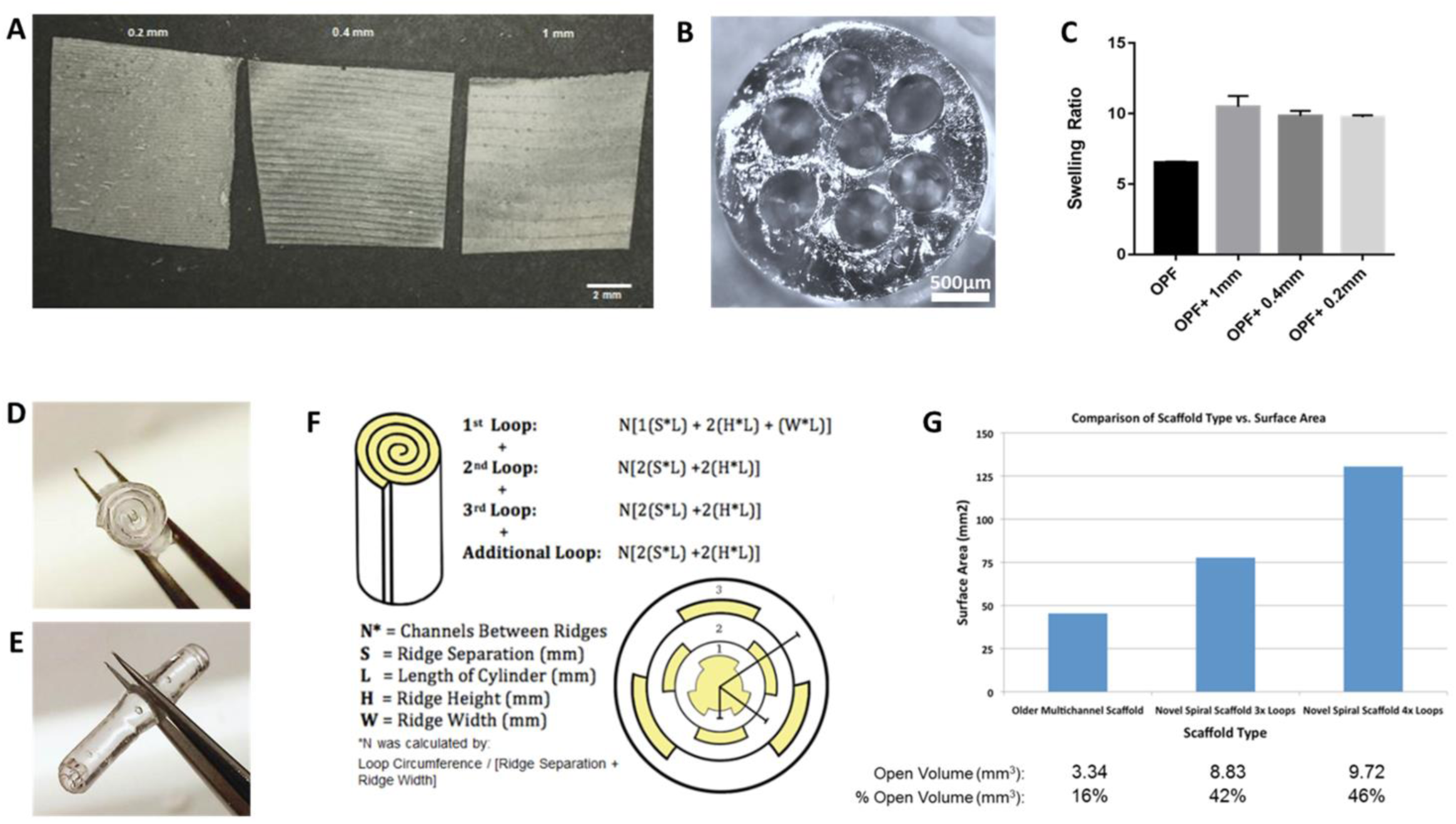
Ridged OPF+ sheet fabrication and characteristics. **(A)** After crosslinking of OPF polymer on Teflon molds, OPF+ sheets contain 100 µm high ridges that are spaced 0.2 mm, 0.4 mm, and 1 mm apart. **(B)** OPF+ has previously been fabricated as a multichannel scaffold with 7 channels and is as wide as a rat spinal cord. We previously demonstrated that axons grown through the channels, as well as the outside surface (Chen *et al*, 2018). **(C)** The swelling ratio of OPF+ sheets were similar to each other and greater than the OPF neutral charged sheet. Bars are represent mean ± SEM. **(D)** and **(E)** When the ridged scaffold is rehydrated it spontaneously rolls up into a 3D configuration along the axis of the ridges. **(F)** Schematic and mathematical modeling demonstrating the calculation of surface area of the scaffold ridged sheet. **(G)** The surface area and volumes both show substantial increase compared to the multichannel scaffold configuration. The 1 mm ridged sheets with 3 to 4 loops maximize the area available for growth.

### 3.2 Whole DRG neurite outgrowth is enhanced on laminin-coated sheets

Whole DRGs were used to determine the effect of different ECM protein coatings (laminin, fibronectin, and collagen) on promoting neurite outgrowth on OPF+ sheets with ridges 1 mm apart. The explants were observed to consistently settle on or immediately adjacent to a ridge, rather than on the broader, flat intervening surfaces (Figure 2). The DRGs elongated along the ridge, and into the adjacent areas in conditions where ECM protein coatings were present in contrast to OPF+ sheets with serum coating. Within 24 hours of culture, laminin-coated OPF+ sheets had longer neurite outgrowth (708.5 ± 34.93 µm, Figure 2E) than controls with serum coating only (255.7 ± 37.12 µm, p=0.004), fibronectin (288.3 ± 33.72 µm, p<0.0001) and collagen (433.6 ± 74.56 µm, p=0.0362). Outgrowth from collagen and fibronectin was not significantly different from serum coated control surfaces. After 48 hours of culture, laminin coated OPF+ sheets (1261 ± 58.98 µm, p <0.0001) continued to support greater neurite outgrowth than serum coated control (445.1 ± 61.15 µm), fibronectin (731.9 ± 69.99 µm, p<0.0001), and collagen (817.3 ± 89.9 µm, p<0.0001), with collagen and fibronectin not significantly different to serum coated control.

**Figure 2:**
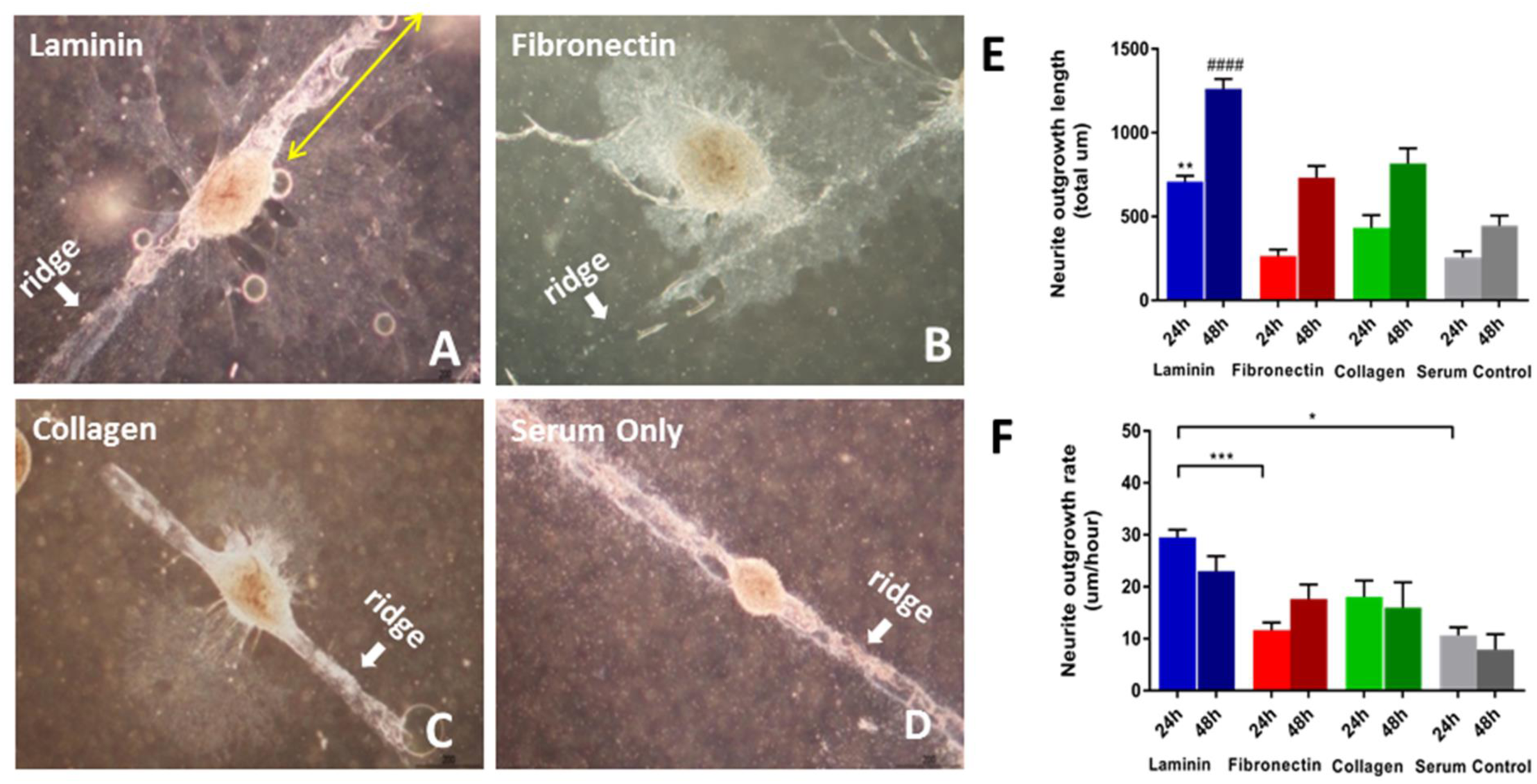
Whole DRG explants on 1 mm spaced OPF+ sheets 48 hours after culture. DRGs were grown on sheets coated with **(A)** laminin, **(B)** fibronectin, **(C)** Collagen, or **(D)** serum only. It was observed that the explants like to attach near or on the ridges and neurites can be observes aligning on the ridge. The long neurite length was measured (yellow arrow) **(E)** Neurite length of DRG explants on differently coated OPF+ sheets 24 and 48 hours after culture (* compared to 24 hours on serum coated sheets; # compared to 48 hours on serum coated sheets). Laminin coated sheets displayed significantly longer neurite outgrowth of DRG explants after 24h and 48h of culture. **(F)** Neurite outgrowth rates in the first and second 24 hours after culture on differently coated 1mm spaced OPF+ sheets. Neurite outgrowth rate in the first 24h of culture of DRG explants on laminin coated sheets were significantly higher than on fibronectin coated and serum only coated sheets. *p<0.05, **p<0.01, ***p< 0.001, ****p<0.0001. Data represent means ± SEM.

The outgrowth rate (µm/h) of whole DRG neurites over 48 hours in culture between the different ECM coatings (Figure 2F) was compared. In the first 24 hours laminin (29.52 ± 1.45 µm/h, p=0.017) enabled a higher rate of neurite outgrowth than was observed on OPF+ sheets with serum coating (10.66 ± 1.55 µm/h). There was no significant difference between the outgrowth rates of fibronectin (11.64 ± 1.46µm; p=0.9977) and collagen (18.07 ± 3.11 µm/h; p=0.9065) compared to serum coating. In the next 24 hours there were no significant differences between laminin (23.03 ± 2.85 µm/h; p=0.1120), fibronectin (17.64 ± 2.8 µm/h; p=0.6721), and collagen (15.99 ± 4.87 µm/h; p=0.8394) as compared with serum coating (7.88 ± 2.98 µm/h).

### 3.3 Dissociated DRG neuron neurite outgrowth is enhanced on laminin coated sheets

OPF+ sheets with ridges 1 mm apart and laminin, fibronectin or collagen ECM supported dispersed neuronal cell attachment and outgrowth. On all surfaces, dissociated neurons preferentially attached to the ridges, and aligned their axons longitudinally along the ridge (Figure 3, Supplemental Figure 1). The use of laminin, fibronectin, or collagen increased the number of adherent neurons on the ridge by 3.4 ± 0.62 fold, 2.69 ± 0.50 fold, and 3.7 ± 0.32 fold, respectively over ridged sheets with serum coating (Figure 3E). The use of laminin resulted in a greater increase in neuronal attachment (38.1 ± 5.98 fold) in the spaces between the ridges compared to collagen coating (4.25 ± 3.3 fold) (p=0.003; figure 3F). Neuronal attachment in between the ridges on fibronectin-coated sheets (32.36 ± 5.65 fold, p=0.013) was also greater statistically than those attached on collagen-coated sheets. 71.67 ± 10.76% (p=0.0127) of neurons attached to the ridges on laminin coated sheets, 60.33 ± 10.67% (p=0.0374) on fibronectin coated sheets, 98.33 ± 12.32 % (p<0.0001) on collagen coated sheets, and 94.33 ± 12.32% (p<0.0001; 2-way ANOVA/Sidak’s multiple comparison) on serum coated sheets, demonstrating a preferential neuronal attachment onto the ridge (figure 3G).

**Figure 3:**
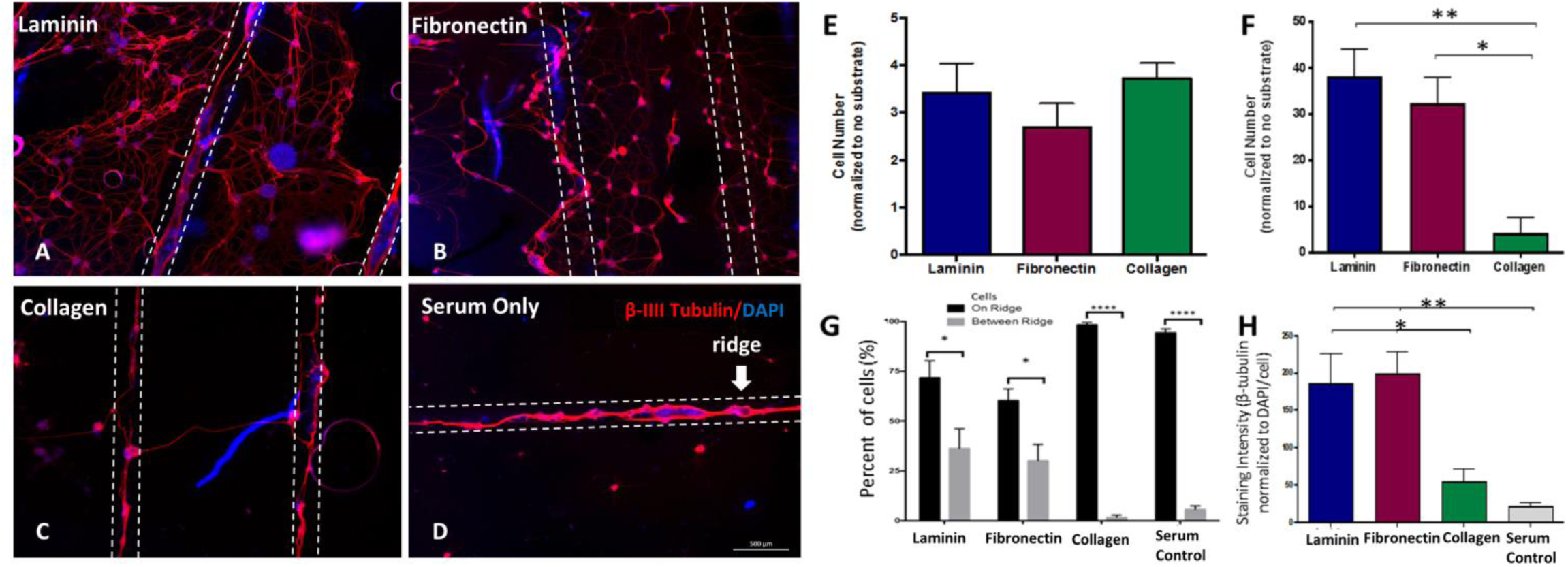
Attachment, alignment, and outgrowth of dissociated DRG neurons on coated OPF+ sheets with ridges 1 mm apart. Immunofluorescent images of b-III tubulin stained axons show attachment and outgrowth from neurons near or on the ridges on OPF+ sheets coated with **(A)** laminin, **(B)** fibronectin, **(C)** collagen, or **(D)** serum only. The neurites preferentially aligned on the ridges (located between the dotted white lines). The neurons are stained with β-III-Tubulin (red) with a DAPI (blue) counter stain. **(E)** The number of neurons on ridges (normalized to the number of cells on the ridges with no substrate) of OPF+ sheets coated with laminin, fibronectin, and collagen. **(F)** The number of cells in between the ridges (when normalized to OPF+ sheets with serum only). **(G)** Percent of cells on ridges or in between ridges for each condition. **(H)** Neurite density as measured by staining intensity (mean gray value) of β-III-Tubulin normalized to the mean gray value of DAPI/cell. *\*p<0.05, **p<0.01, ***p< 0.001, ****p<0.0001. Data represent means ± SEM.*

The neurite density of dissociated neurons was also greater on laminin and fibronectin coated sheets (figure 3H), as measured by stereologic analysis of the areas of β-tubulin staining. Laminin coated sheets (185.3± 41.02 mean gray value; p=0.002) and fibronectin (198.4± 30.14 mean gray value; p=0.0408) had higher total β-tubulin staining per cell than sheets with serum coating (20.73 ± 5.59 mean gray value). The neurite density on laminin coated sheets (p= 0.0263) and fibronectin coated sheets (p=0.0408) was also greater than that seen on collagen coated sheets.

### 3.4 Increasing the number of ridges improves neuronal cell attachment, alignment, and neurite density

To determine what effect ridge spacing distance may have on neuronal cell attachment and alignment, laminin-coated OPF+ sheets with ridges 0.2 mm, 0.4 mm, and 1 mm apart were used. Tighter ridge spacing increased the number of ridges per area and the overall surface area of the sheet, (figure 4 A-C) which in turn improved neuronal cell attachment. Many of the cells settled around or on top of the ridges, extending neurites along the ridge. The number of neuronal cells located on the ridge surface was 2.5 fold greater for the 0.2 mm spaced OPF+ sheets (figure 4 D; 208.6 ± 44.3 cells; p=0.039) than the 1 mm OPF+ sheets (84.8 ± 15.1 cells). The 0.4 mm OPF+ sheets (figure 4 D; 186.4 ± 11.9 cells; p=0.1215) had 2 fold greater number of cells than the 1 mm spaced OPF+ sheets. The neurite density was increased on the OPF+ sheets with ridges more closely spaced (0.2 mm and 0.4 mm apart). We found a 2.9 fold increase in the amount of β-III tubulin staining (normalized to the amount of DAPI staining /cell) when comparing OPF+ sheets with 0.2 mm spaced ridges (figure 4 E; 719.5 ± 130.3 mean grey value; p=0.012) to OPF+ sheets with 1 mm spaced ridges (251.2 ± 33.4 mean grey value).

**Figure 4:**
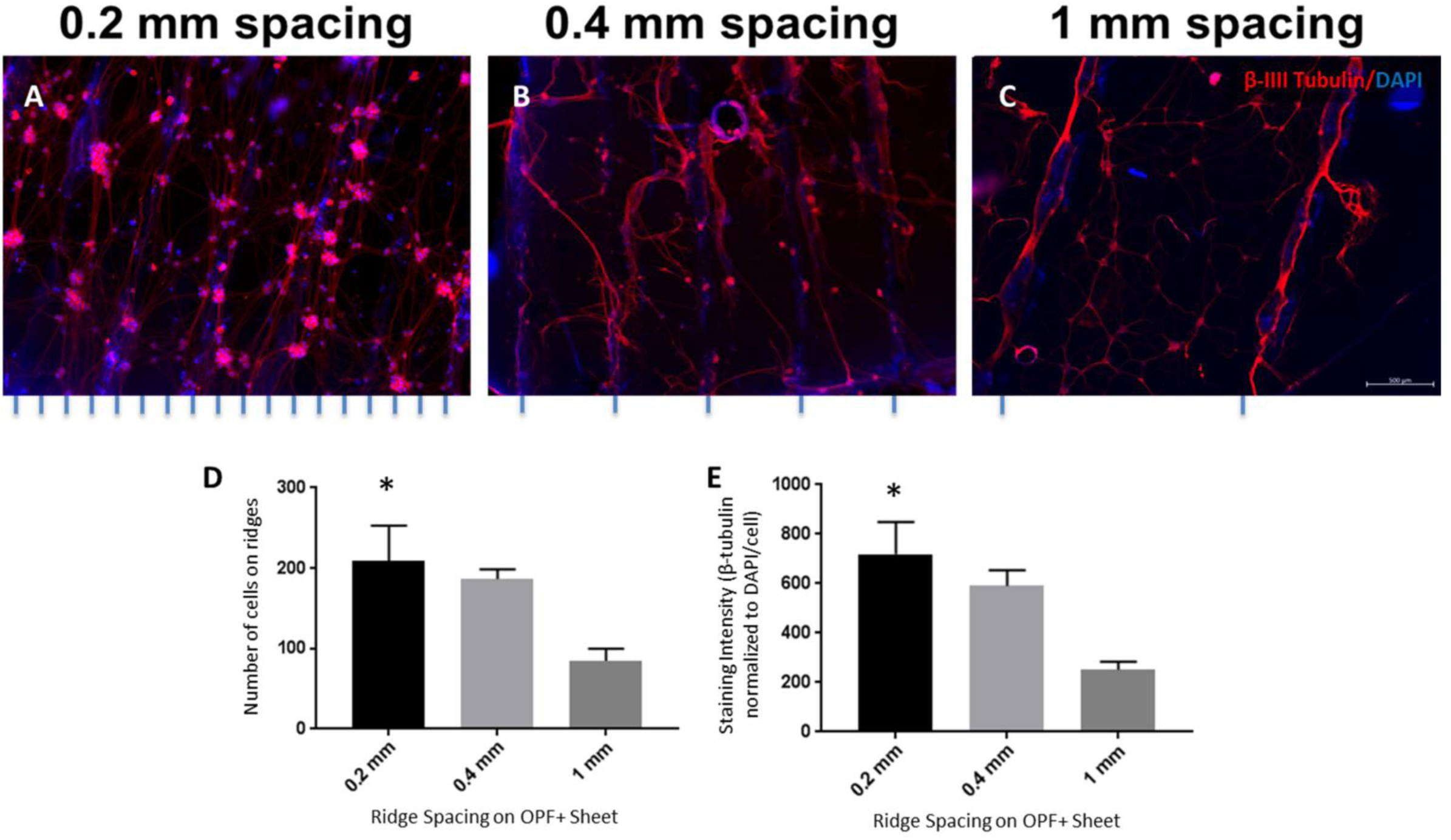
Alignment, attachment, and neurite density of disassociated DRG neurons on OPF+ sheets with ridges spaced 0.2 mm, 0.4 mm, and 1 mm. The neurons were labelled using β-III tubulin (red) and DAPI (blue). When the ridges are spaced closer together, such as 0.2 mm apart **(A)** or 0.4 mm apart **(B)**, there are more neurites observed aligned along the ridges than the 1 mm spaced sheets **(C). (D)** Number of cells on the ridges of the OPF+ scaffolds. **(E)** β-III tubulin staining intensity (normalized to DAPI/cell) of neurons grown on OPF+ sheets with ridges spaced 0.2 mm, 0.4 mm, and 1 mm apart. *\*p<0.05 compared to 1mm spacing as control. Data represent means ± SEM.*

### 3.5 The number of ridges influences Schwann cell organization and axonal myelination

In our previous studies we have loaded the multichannel scaffolds with genetically modified Schwann cells which facilitated axonal regeneration and recovery after spinal cord transection [5]. We next sought to demonstrate that the OPF+ ridged sheet design with laminin coating would support the growth of Schwann cells. Schwann cells on OPF+ ridged sheets initially formed a cell layer in a dispersed and unorganized manner (Figure 5 A-C). Over a 6 day period the Schwann cells organized along the ridges of the scaffold (Figure 5 D-F). When the Schwann cells were co-cultured with dispersed DRG neurons for 4 weeks, neurites became myelinated (Figure 6). Neurites aligned well with the ridges and stained positively for myelin basic protein (MBP). Greater densities of aligned myelinated neurites were observed when the ridges were spaced closer together (0.2 mm spacing vs. 1 mm spacing).

**Figure 5:**
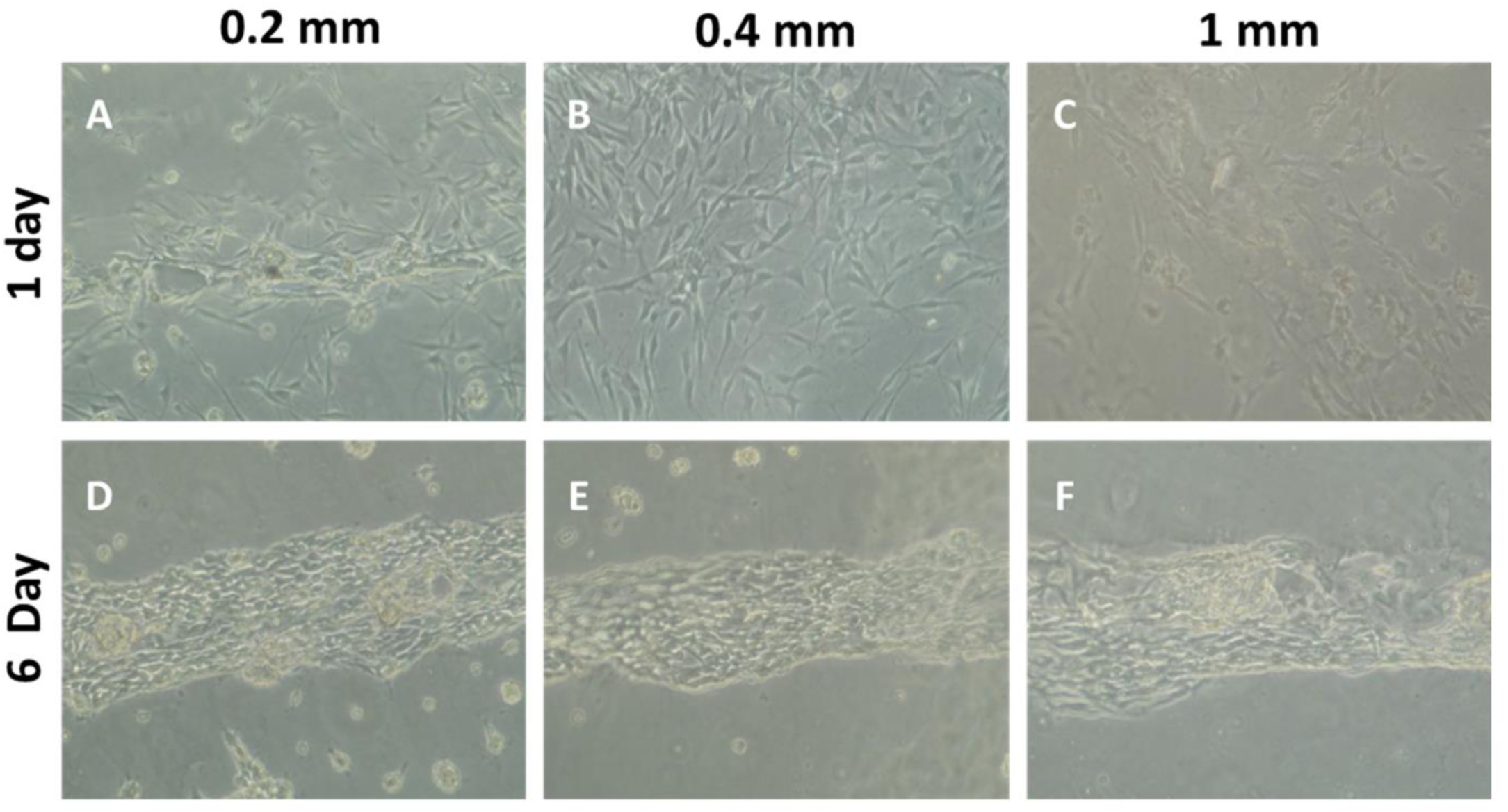
Bright field images of primary rat Schwann cells grown on OPF+ sheets with ridges 0.2 mm, 0.4 mm, and 1 mm apart. One day after culture the Schwann cells cover the sheet in an unorganized manner **(A-C)**. However, 6 days after culture the Schwann cells are aligned along the ridges **(D-F)**.

**Figure 6:**
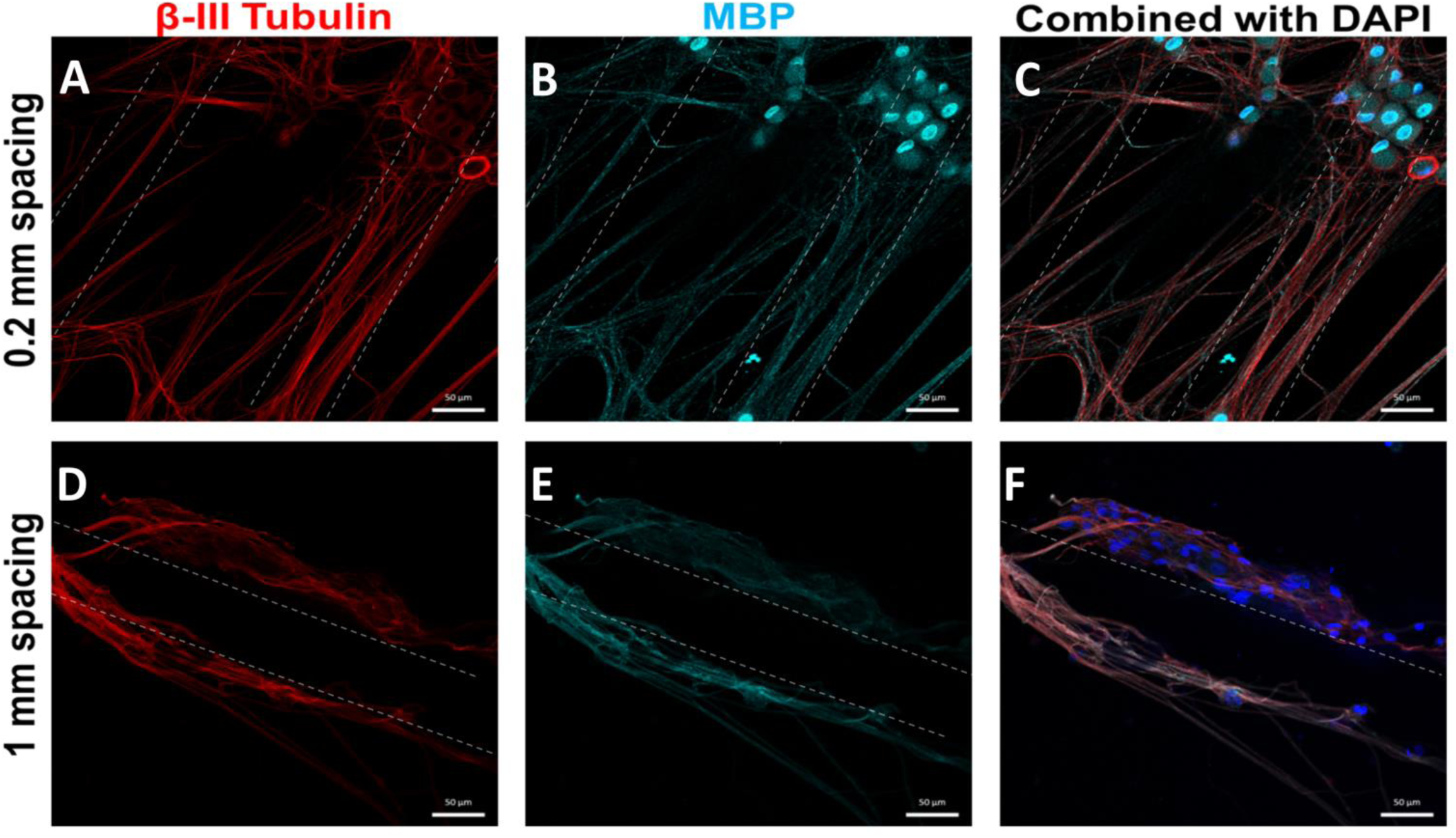
Confocal images showing Schwann cells co-cultured with disassociated DRG neuron on OPF+ sheets with ridges spaced 0.2 mm or 1 mm apart. The cells were labelled with β-III tubulin staining (red, **A, C, D & F**), Myelin Basic Protein (teal, **B, C, E & F**), and DAPI (blue, **C & F**). After culture in media that supports myelination by Schwann cells, the Schwann cells myelinate the disassociated DRG neurites. Since alignment is enhanced by having the ridges on the OPF+ sheet closer together, it can be observed that there are more aligned myelinated neurites on the 0.2 mm spaced OPF+ sheets **(A-C)** than 1 mm spaced sheets **(D-E)**.

### 3.6 Rolled OPF+ scaffold sheets, housed within a single channel scaffold, integrated into the spinal cord and supported axon outgrowth into the scaffold after complete transection injury

A proof of principle pilot experiment was conducted in which uncoated and laminin-coated OPF+ sheets with ridges 0.4 mm apart, were rolled and secured inside a single channel OPF+ scaffold (Figure 7A). This scaffold was then implanted into the transected rat spinal cord at the T9 level. The scaffold integrated well with the spinal cord over a 4 week period (Figure 7B). Staining for β-III tubulin indicates that both scaffolds promoted axon outgrowth into the scaffold (Figure 7C-D). Although the number of animals was not powered to determine statistical significance (n=1 per condition), these preliminary results show the potential of our novel rolled scaffold for spinal cord regeneration.

**Figure 7:**
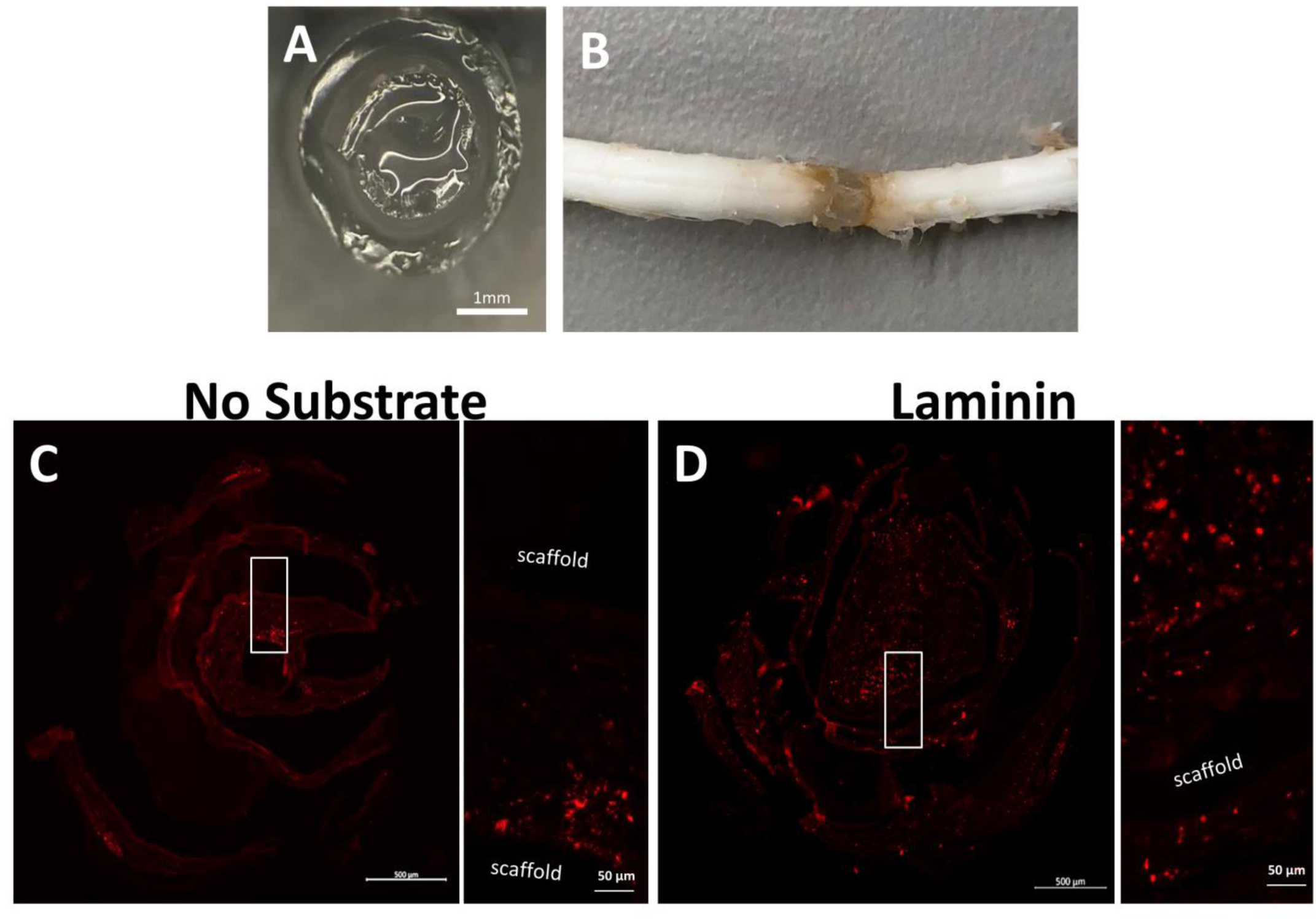
Proof of principle implantation of rolled OPF+ sheets with ridges spaced 0.4 mm apart for 4 weeks following transection and implantation. **(A)** Rolled OPF+ sheet with ridges spaced 0.4 mm apart was coated with laminin or no substrate and then placed in a single channel OPF+ scaffold to hold the structure together. **(B)** The OPF+ scaffold was implanted into a transected T9 spinal cord and demonstrated good integration into the spinal cord 4 weeks post implantation. **(C & D)** Cross section of scaffolds containing regenerating axons labelled by neurofilament antibody 4 weeks after implantation.

## 4. Discussion

We have previously demonstrated that OPF+ fabricated in a multichannel design with seven channels can enhance regeneration after SCI [4, 7] in rats. This regeneration can be further improved by loading the channels with GDNF-secreting Schwann cells, leading to modest functional recovery [5]. We have also shown that the OPF+ material can be modified by incorporating the anti-fibrotic drug rapamycin in PLGA microspheres [27]. OPF+ multichannel scaffolds containing rapamycin microspheres were also combined with Schwann cells to help promote functional recovery following SCI. It was also noted from these studies that many of the regenerating axons grew on the outer circumferential surface of the scaffold. We therefore hypothesized that axonal growth may preferentially occur on surfaces rather than within open spaces. Maximizing the surface areas available to regenerating axons to extend may further improve the density and directionality of regrowth, and impact upon functional reconnections. In this study, we have now demonstrated, *in vitro*, that OPF+ fabricated in a flat sheet design with ridges increased the surface area available for growth when compared to the multichannel design scaffold. Providing an attractive ECM protein coat and spacing the ridges closer together improved cell attachment, outgrowth, and alignment of DRG explants, neurons, and Schwann cells. We also demonstrated that this scaffold design is feasible to use *in vivo*.

When hydrated, the OPF+ sheets spontaneously form a 3D spiral tube. Hydrogels have been designed to self-roll in previous studies either by introducing reductants to cleave disulfide bonds [28], changes in pH and sacrificial layers [29], or using light responsive inks [30] and photolithography [31]. These ridged scaffolds had swelling ratios comparable to our previous studies [32]. Hydrogels may suffer from low cell attachment due to the hydrated surface layer [33]. This issue could be addressed by coating the hydrogel sheets with collagen, laminin, or fibronectin, all of which are classically established as ECM coatings used to promote neurite outgrowth [6, 13, 34, 35]. The current study shows that DRG explants cultured on laminin coated OPF+ sheets had longer neurite outgrowth than fibronectin or collagen coated sheets. In addition, the outgrowth rate of neurites on laminin coated sheets was faster in the first 24 hours of culture. This can be due to the different amount of proteins used or absorbed into the scaffold. Previous studies have demonstrated concentration dependent differences in cell attachment and outgrowth, however, concentrations used in this study were within range of those used in other studies [6, 24, 36, 37]. It is also interesting to note that the DRG outgrowth rate on the fibronectin coated sheets was still increasing at the 24-48 hour time point and may have results in greater outgrowth length at a later time point. When neuronal attachment was measured on or in between the ridges, laminin and fibronectin did equally well. The observed enhancement of neuronal outgrowth and guidance associated with laminin is in keeping with several other studies investigating different substrates for neuronal regeneration [6, 13, 35, 38]. ECM proteins or their fragments play an important role in cell adhesion through ligand receptor interactions, and these interactions may be an important component to consider in developing novel biomaterials [39-41]. In addition, the three dimensional structure of ECMs help anchor cells and provide axonal guidance [15]. We show that to maximize the ability of hydrogel scaffolds to anchor neurons and their axons and aid in their outgrowth, ECMs like laminin and fibronectin are necessary to maximize the regenerative capability of the scaffold.

The combination of a scaffold containing ridges and ECM may mimic what is seen in development where topographical and chemical cues help guide axons. Factors such as stiffness of the substrate can be important in determining the growth patterns of neurons, as shown by the growth of retinal ganglion cell axons towards softer tissue [42]. Laminin and β-integrin signaling have been found to be important in neuronal polarization and the deletion of β-integrin results in deficits in axonal development [43]. Radial glia, similar to the ridges of the scaffold, provide topographical cues that orientate the migration of embryonic neurons [44]. Neurite outgrowth from DRG explants and disassociated DRG neurons could be directed in a bidirectional way on ridged surfaces in this study. Ridges may provide important topographical cues, as the majority of neurons preferentially attached on or near the ridges of the scaffold. A similar preference for mechanical guidance cues (microgrooves), rather than on flat spaces between the microgrooves, has previously been observed [13]. Studies investigating the effect of grooves on a polymer surface have shown that axonal alignment and behavior could be influenced by the depth of the grooves, spacing between the grooves, and groove widths [13, 38]. Similarly, in our study, disassociated DRG neurons that grew on the OPF+ sheets with ridges spaced 0.2 mm apart had greater cell attachment to the ridges, alignment, and outgrowth than those grown on 1 mm spaced sheets. One of the goals of the scaffold is to bridge the gap that is created after spinal cord injury. The ability of the scaffold to direct growth in a longitudinal direction and maximize this directed growth is an important consideration.

Another approach to enhancing regeneration is to mimic the peripheral nervous system environment. It is well established that the peripheral nervous system has a better ability to regenerate than the central nervous system. This may be due to a more rapid clearance and recycling of myelin debris, more permissive ECM, and a neurotrophic Schwann cell response [45]. The use of Schwann cells with biomaterials recreates a peripheral nervous system environment in the spinal cord, where the cells can contribute regenerative ECM, growth factors, and facilitate remyelination. Our lab, among others, has compared different cell types. Schwann cells have been found to be best able to support axonal regeneration [10, 26, 46, 47]. In this study, Schwann cells preferentially organized along the ridges of OPF+ sheets and myelinated neurites. The OPF+ sheet with ridges spaced 0.2 mm apart had more aligned myelinated neurites observable per area than the 1 mm sheet. This combination of the OPF+ ridged sheet with an ECM and Schwann cells may increase the regenerative potential of the scaffold and these *in vitro* results suggest that this combination would be helpful in guiding and myelinating aligned axons.

This study demonstrates a novel hybrid design of the OPF+ scaffold for use in regenerative medicine, particularly suited for spinal cord injury. It has properties similar to the spinal cord, maximizes available surface area and volume for regeneration, and can spontaneously roll to make a 3D structure for transplantation. Flattened, ridged sheets were used for *in vitro* studies and demonstrated improved neuron attachment, axonal alignment, and outgrowth. Furthermore, it would be straightforward to manufacture and scale for large animal studies and clinical applications. Scaffold properties can be modified including varying the ridge distance and size, as well as, coating with different substances. It is also possible to study different cell types for different applications in regenerative medicine. In a small study when implanted into transected spinal cord, the rolled scaffold integrated well and supported regenerating axons. This scaffold is well positioned to be used for combinatorial treatments to treat spinal cord injury through use of novel biomaterials, ECMs, and cells.

## Acknowledgements

The authors would like to acknowledge Jewel Podratz with training and assistance with primary cell culture of DRGs and Ann Schmeichel with training and assistance with immunocytochemistry techniques. We would also like to acknowledge Bingkun Chen and Jarred Nesbitt for their assistance with the animal surgery and post-surgical care. We also thank Shuya Zhang for her help with sectioning of the spinal cord tissue.

## Conflict of Interest

None.

---

**Supplemental Figure 1:**
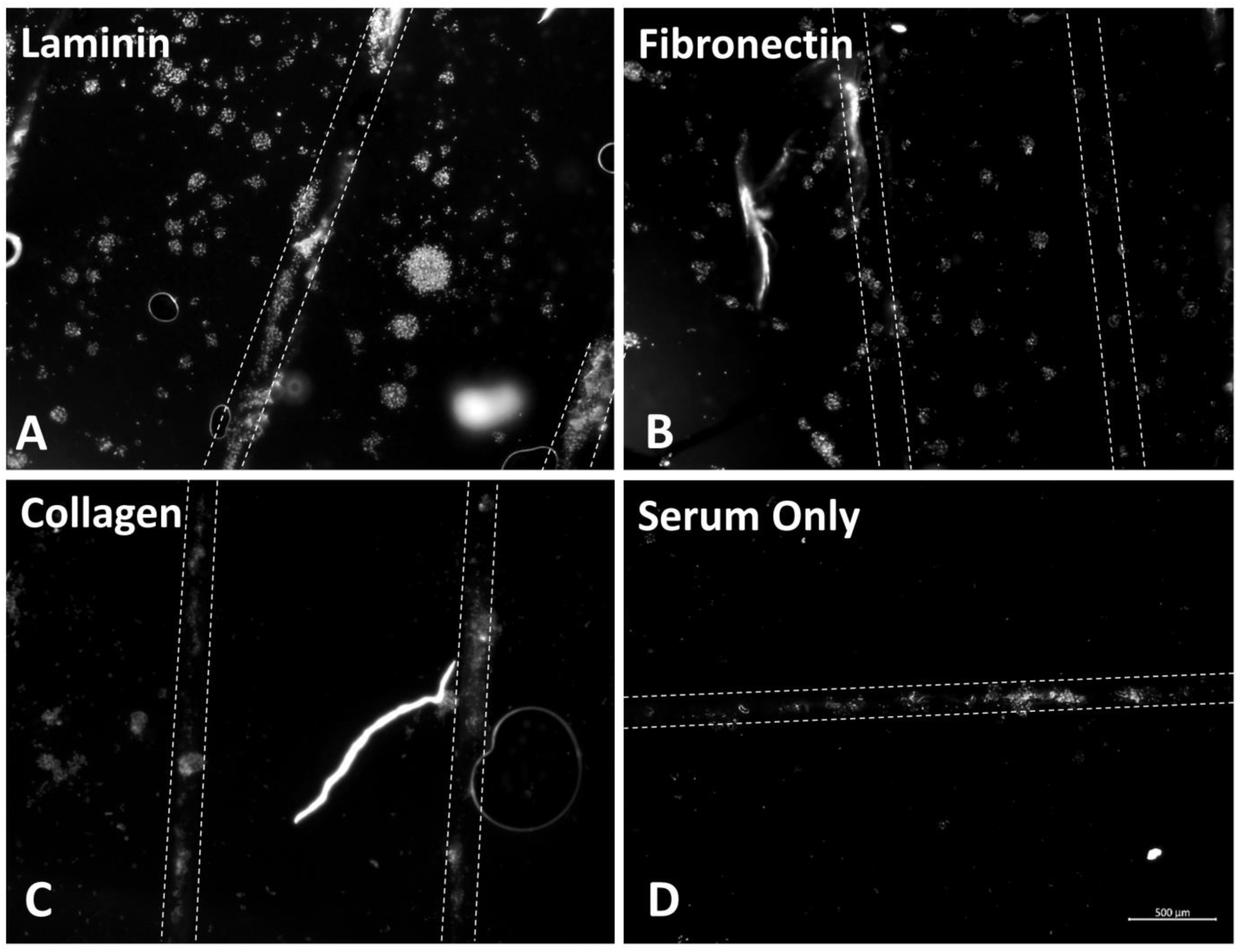
DAPI staining of dissociated DRG neurons on OPF+ scaffold sheets with ridges 1 mm apart. The sheets were coated with (A) Laminin, (B) Fibronectin, (C) Collagen, and (D) Serum only. Number of cells on and off ridges was determine by DAPI co-labelled with b-III tubulin (Figure 3).

## Notes

Grant Support: This publication was supported by Grants from the New Jersey Commission on Spinal Cord Research CSCR15IRG002 and the National Center for Advancing Translational Sciences (NCATS) UL1 TR002377 and TL1 TR002380. It was also supported by grants from Regenerative Medicine Minnesota and the Bowen Foundation, the Kipnis Foundation and the Mayo Clinic Center for Regenerative Medicine.

